# A phylogenetic method linking nucleotide substitution rates to rates of continuous trait evolution

**DOI:** 10.1101/2023.10.04.560937

**Authors:** Patrick Gemmell, Timothy B. Sackton, Scott V. Edwards, Jun S. Liu

## Abstract

Genomes contain conserved non-coding sequences that perform important biological functions, such as gene regulation. We present a phylogenetic method, PhyloAcc-C, that associates nucleotide substitution rates with changes in a continuous trait of interest. The method takes as input a multiple sequence alignment of conserved elements, continuous trait data observed in extant species, and a background phylogeny and substitution process. Gibbs sampling is used to assign rate categories (background, conserved, accelerated) to lineages and explore whether the assigned rate categories are associated with increases or decreases in the rate of trait evolution. We test our method using simulations and then illustrate its application using mammalian body size and lifespan data previously analyzed with respect to protein coding genes. Like other studies, we find processes such as tumor suppression, telomere maintenance, and p53 regulation to be related to changes in longevity and body size. In addition, we also find that skeletal genes, and developmental processes, such as sprouting angiogenesis, are relevant. The R/C++ software package implementing our method is available under an open source license from https://github.com/phyloacc/PhyloAcc-C.

## 1 Introduction

In recent years there have been numerous advances in mapping genes underlying phenotypic traits. Many of these advances have built on the successes and refinements of traditional genetic mapping methods yet, as is articulated by Smith et al. [1], such approaches are often limited to the small number of model organisms amenable to crosses or other genetic manipulations. Recently, alternative phylogenetic approaches driven by comparative genomics have emerged as a useful tool for mapping genes in species not amenable to traditional approaches. Several methodologies have been proposed to associate evolution of genes or genomic regions with changes in phenotypic traits including those of [2, 3, 4, 5, 6, 7]. These studies use a variety of genomic signatures as evidence of association with phenotypic evolution, including increases in evolutionary rate, loss of function such as pseudogenization, or wholesale deletion of genes or non-coding regions from the genome. Comparative approaches of this kind (hereafter ‘PhyloG2P’) have proved to be surprisingly powerful at identifying associations between genomic and phenotypic variation in the context of convergent evolution of the phenotypic trait.

With the PhyloG2P research programme in mind, this paper aims to make three contributions. First, we highlight the idea of relating phenotypic and genotypic evolution by linking substitution rate multipliers (for nucleotide changes) to variance multipliers (for changes in a continuous trait). Second, we introduce a specific piece of software, PhyloAcc-C, that applies this approach in the context of conserved non-coding elements (CNEs). Third, we illustrate the PhyloAcc-C software using real data, running it with a set of mammalian CNEs and a lifespan related trait as input, thereby providing an opportunity to discuss its output in the context of other recent PhyloG2P-style studies.

The overarching biological motivation for our study is our interest in evolutionary innovation, and in this paper we are particularly concerned with methods that attempt to answer the question ‘which CNEs are related to changes in a continuous trait I care about?’ This is a salient question as it has been recognized for decades that there are many highly conserved stretches of non-coding DNA that participate in gene regulation across diverse species e.g. [8]. Indeed, such sequences are routinely annotated [9] in the UCSC Genome Browser [10] and may then be related to the evolution of phenotypic traits. To give one example, Booker et al. [11] identified conserved sequences that were accelerated specifically in bats, and showed that a subset acted as limb enhancers in transgenic mice. Their conclusion was that some identified enhancers were potentially instrumental to the evolution of bat wings. This conclusion was reached without modelling the co-variation of the rate of enhancer evolution and key measurements from bat wings. However, it is possible that incorporating measurements such as limb length could highlight additional relevant loci that had experienced more subtle evolutionary trajectories than bat specific acceleration.

More generally, as reviewed by Smith et al. [1], there is widespread interest in relating phenotypic and molecular evolution. This interest is evidenced by the substantial effort put into producing a variety of software packages and studies that quantify the relationship between traits and substitution rates. Examples include Forward Genomics [4] and reverse genomics [3], both of which relate sequence similarity (via correlation, generalized least-squares, or heuristics) to traits (with ancestral values inferred using parsimony algorithms). A methodology with a similar goal is that of Treaster et al. [7], which uses tree topology to model the intuitive notion that comparisons between more closely related species should be less confounded by genetic background than comparisons between more distantly related ones. A recent contribution to this diverse collection of approaches is PhyloAcc [12], a Bayesian phylogenetic approach centered on latent conservation states, which is modified here in this paper. A key feature of the above four approaches is that they deal with discrete traits, and in the case of PhyloAcc, do not explicitly model the trait, instead relying on a priori reconstruction, often under the assumption of convergent gain or loss of a character state.

Methods for studying the relationship between continuous traits and molecular evolution are fewer. The Coevol approach [13] models the co-evolution of continuous traits and rates using a multivariate diffusion process, and does not require user supplied branch lengths, though calibrations can be supplied if desired. One imagines that constraining branching times will sometimes be helpful, especially when the sequences being considered are short and highly conserved, and therefore contain few distinguishing differences, as is the case with mammalian CNEs. Two more empirically focused recent studies are that of Yusuf et al. [14], who study the co-evolution of bill shape and both protein coding and non-coding DNA, and that of Kowalczyk et al. [15], who study the lifespan and body size of mammals. The former study used k-means binning to group branches of a tree based on the rate of trait evolution, and then used a likelihood ratio test to compare nucleotide substitution rates under a global clock model versus a local clock model, with one rate per bin. The latter study used the RERConverge method [6], which correlates relative rates of protein evolution and ancestral state reconstructions of a continuous trait, each estimated separately using maximum likelihood.

Here we describe the PhyloAcc-C model, which connects the evolution of continuous traits and non-coding DNA using a statistically integrated approach. We then illustrate the use of our model by applying it to a mammalian trait previously analyzed using RERConverge, but this time considering CNEs rather than protein coding genes, thereby providing analyses that complement the existing literature.

## 2 Methods

The PhyloAcc-C method follows the general Bayesian approach taken by Hu et al. [12] and modifies it so as to model continuous phenotypic change. In this section, we describe the method in enough detail that it may be recreated, and so that one may understand or modify our open source R/C++ implementation.

### Input

The method relies on four inputs: (1) a rooted phylogeny **T** having *L* leaves, *N* = 2*L −* 1 nodes, and *E* = *N −* 1 edges, and that encapsulates the relationships between species from which trait data is drawn; (2) a multiple sequence alignment **X**_*L×S*_ juxtaposing homologous CNE sequences observed in the *L* species (rows) at *S* sites (columns); (3) a vector **y** = (*y*_1_, *y*_2_, …, *y*_*L*_) of continuous trait measurements observed in the corresponding species; (4) a rate matrix **Q**_4*×*4_ and stationary distribution *π* that model the background nucleotide substitution process at putatively neutral sites. The alignment may contain gaps which will be treated as missing data. Both the alignment and the nucleotide substitution parameters can be obtained using standard methods e.g. as detailed by Hu et al. [12]. In particular, the rate matrix is often estimated using methods like PhyloP [16], which can estimate **Q** from an alignment of putatively neutral and easily alignable sites, such as the third position of codons.

### Model

Each branch is assigned a conservation state *z*_*i*_ that takes on three values: background (*z*_*i*_ = 1), conserved (*z*_*i*_ = 2), or accelerated (*z*_*i*_ = 3). This categorization of branches into three states follows that of Hu et al. [12], which in turn was based on the approach taken by Pollard et al. [16], who apply a conserved and an accelerated state to branches of a tree in the PhyloP software. Conservation states are not assigned freely but follow a Markov process from root to tips so that *p*(*z*_*j*_|*z*_*i*_) = **Φ**_*i,j*_ where

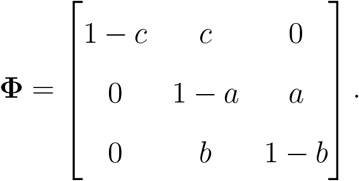

The structure of this matrix allows CNEs to become conserved and later accelerated. Because a transition from accelerated back to conserved is possible (i.e *b >* 0), bursts of acceleration can occur on internal branches. In principle, other matrices can be used to either constrain or relax the transition between rate categories across the tree.

The conservation state of a branch affects both the nucleotide substitution process and the rate at which a trait evolves. Conservation states modulate nucleotide substitution rates via substitution rate multipliers **r** = (*r*_1_ = 1, *r*_2_, *r*_3_) so that the probability of transition from nucleotide *a* to *b* on branch *i* of length *t*_*i*_ is expm*{***Q** *• r*_*z*_*i • t*_*i*_*}*_*a,b*_.

Similarly, conservation states on branches modulate the magnitude of trait changes along branches via variance multipliers **v** = (*σ*^2^, *β*_2_*σ*^2^, *β*_3_*σ*^2^). Under the model, traits evolve according to normally distributed displacements along the branches of **T** such that cumulative displacements are observed in *y*_1_, …, *y*_*L*_. Displacements have mean 0 and a variance that is proportional to both the branch length *t*_*j*_ and the appropriate variance multiplier:

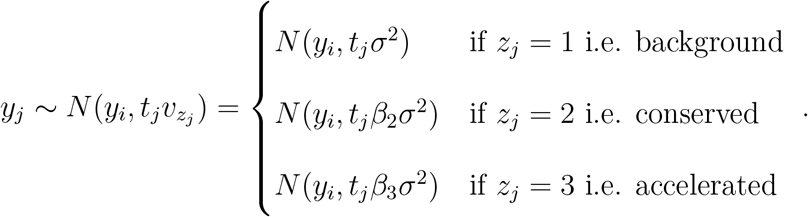

### Joint likelihood

Letting *pa*(*i*) denote the parent of node *i*, and assuming that *pa*(*j*) = *pa*(*k*) = *i*, we can write the joint likelihood of the model as:

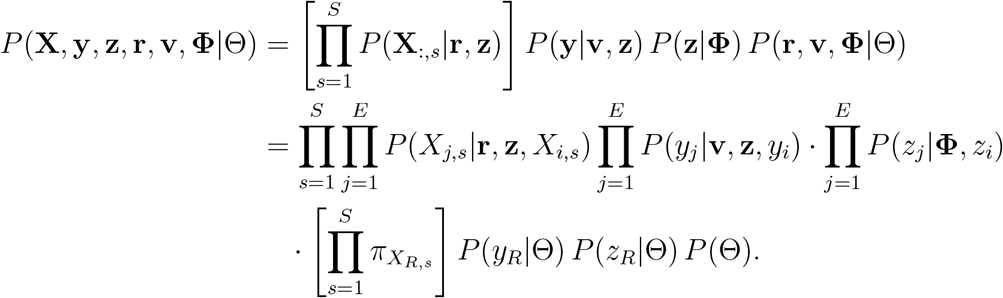

The last line of the above product indicates that at the root node *R* the prior probability of observing a nucleotide is given by the (input) stationary distribution *π*, whereas the trait and conservation state are specified directly using a prior, as described below.

### Specification of priors

At the root node, we use *p*(*z*_*R*_ = 1) = *p*(*z*_*R*_ = 2) = 0.5 and *y*_*R*_ *∼ N* (0, 1), although users can make their own choices freely.

Priors on the entries *a, b, c* in **Φ** are uniform distributions (i.e., Beta(1,1)). Because we use the same nucleotide data, *r*_2_ *∼* Gamma(5, 0.04) and *r*_3_ *∼* Gamma(10, 0.2), follow the values in Hu et al. [12].

Priors on log *β*_2_ and log *β*_3_ are *N* (0, 1) which is mathematically equivalent to setting a *N* (0, 2) on log(*β*_3_ : *β*_2_), the logarithm of their ratio. We used a prior of log *σ*^2^ *∼ N* (0, 2).

### Bayesian inference procedure

Inference is performed using a Markov chain Monte Carlo procedure, which is a combination of collapsed Gibbs sampling with some Metropolis within Gibbs steps [17, 18]. The following steps are repeated:

#### Step 1: sample ancestral trait values y and trait variance multipliers v

Perform Metropolis steps (default is 500) to propose and update *σ*^2^, *β*_2_, *β*_3_, and latent **y**. On 60% of iterations proposals to modify *σ*^2^, *β*_2_, and *β*_3_ are made; on the remaining occasions proposals to perturb latent *y*_*L*+1_, …, *y*_*N*_ are made.

#### Step 2: sample ancestral nucleotides X

First use the familiar pruning algorithm [19] to calculate the likelihood of subalignment {**X**_*i,s*_} rooted at node *i* for all sites *s* = 1…*S* using the recurrence:

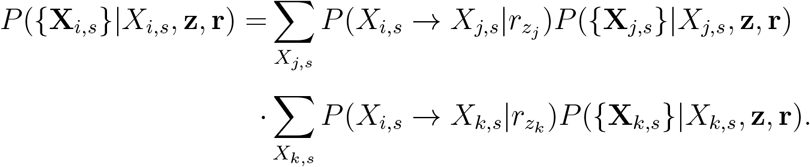

Next, forward sample ancestral nucleotides **X**_*L*+1…*R*,1…*S*_. For sites at the root node we have *P* (*X*_*R,s*_|{**X**_*R,s*_}, **z, r**) ∝ *P* ({**X**_*R,s*_}|*X*_*R,s*_, **z, r**)*P* (*X*_*R,s*_) and 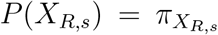 by assumption. For the remaining internal nodes we work from root to tips on a per-site basis using:

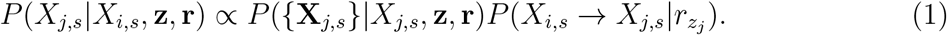

#### Step 3: sample per-branch latent conservation states z

First, from tips to root, calculate the joint likelihood of trait values and nucleotide emissions *{***XY**_*i*_*}* occurring on the subtree rooted at node *i* using the recurrence:

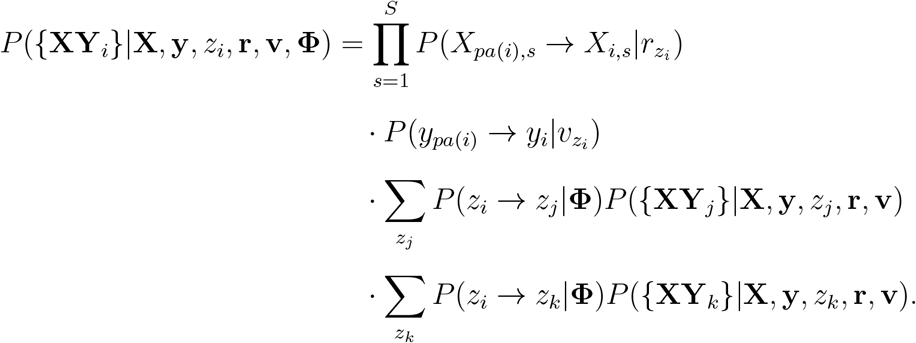

Next, sample **z** from root to tip. At the root our (domain specific) prior is *P* (*z*_*R*_) = (0.5, 0.5, 0.0). For descendant nodes the appropriate probabilities are:

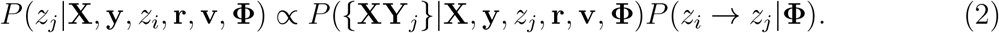

#### Step 4: sample per-category nucleotide substitution rate multipliers r

Perform Metropolis step to propose/update substitution rate multipliers *r*_2_ and *r*_3_.

#### Step 5: sample latent rate category transition probabilities Φ

The beta prior on entries *a, b* and *c* of **Φ** leads to a beta posterior. For example, the posterior of *c* is directly sampled based on **z** transitions from 1 *→* 2 as follows:

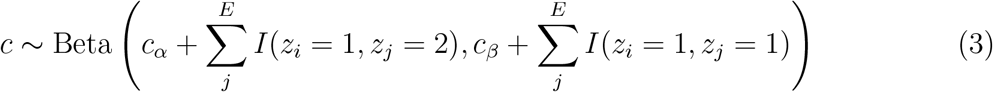

The posteriors of *a* and *b* are calculated similarly using the count of transitions 2 *→* 3 and 3 *→* 2 respectively.

### Model selection and ranking of associated loci

A collection of candidate elements can be ranked for association with a trait of interest using the Bayes factor (BF) in favour of the ‘full model’ described above. In the full model, *σ*^2^, *β*_2_, and *β*_3_ are free to vary whereas in the restricted model this is not the case, and *β*_2_ = *β*_3_ = 1 so that no systematic relationship between the rate of trait evolution and relative substitution rates is specified. As the null hypothesis is nested and the priors on *β*_2_ and *β*_3_ are common to all candidate elements, the BF is estimated using the posterior density of (log*β*_2_, log*β*_3_) at (0, 0). This is an application of the Savage–Dickey method, which is explained in the tutorial of Wagenmakers et al. [20].

### Molecular data

We obtained mammalian CNE alignments directly from the first author of [12], who had in turn originally obtained them from the UCSC 100-way vertebrate alignment [21] available at: http://hgdownload.soe.ucsc.edu/goldenPath/hg38/multiz100way. We obtained the rate matrix and mammal phylogeny (see S1 Text) used to model the background relationship between species from the PhyloAcc GitHub repository, prepared by Hu et al. [12], and available at: https://github.com/phyloacc/Hu-etal-2019-data/. The mammal phylogeny was originally prepared by Murphy et al. [22].

### Software implementation

The PhyloAcc-C software is implemented as an R package [23] that makes use of C++ functions [24, 25] to perform MCMC sampling. To use the package one must load an alignment (e.g. a FASTA file) and a tree (e.g. a New Hampshire file) by using the package ape [26] or similar. Trait data should be loaded into an R data frame and labelled so that it can be matched to the species names in the tree.

The PhyloAcc-C package includes helper functions (sim X, sim y, and sim z) enabling one to simulate alignments, traits, and conservation states under the PhyloAcc-C model. We used these functions to assess model performance in this paper, and a user may do the same given their own phylogeny and rate matrix.

The software, installation instructions, and a tutorial covering simulation and inference are all available at https://github.com/phyloacc/PhyloAcc-C.

## 3 Results

To demonstrate that it is possible in principle to relate the rate of trait evolution to the rate of nucleotide evolution using PhyloAcc-C, we performed a simulation study under ideal circumstances. Then, to illustrate the method using real data, we downloaded principal component data representing the trait ‘long-lived large-bodied’ (LLL) previously studied with respect to protein evolution by Kowalczyk et al. [15]. This trait was analyzed in relation to CNEs previously studied by Hu et al. [12].

In both simulations and our illustrative example, we focused on the quantity log *β*_3_ : *β*_2_. This quantity contrasts variation of a trait on branches undergoing accelerated sequence evolution against variation on branches undergoing conserved sequence evolution; a positive quantity associates faster nucleotide evolution with faster trait change while a negative quantity associates faster nucleotide evolution with slower trait evolution. Values close to zero suggest no strong systematic relationship under the PhyloAcc-C model.

### 3.1 Simulations on full binary trees

Using a fully bifurcating ultrametric tree with 128 tips, we simulated 100 times from our prior distribution, generating latent conservation states, and corresponding DNA sequence alignments and phenotypic trait values. The length of the simulated elements was 80 bp, reflecting the median length of mammalian CNEs in our data set. Branch lengths were all set at 0.1, so that root to tip distances were similar to the longer of the root to tip distances on our mammalian tree.

We were able to recover log *β*_3_ : *β*_2_ reasonably well, with a MSE (mean squared error) of 0.36 (Figure 1A). The estimates appeared reasonably calibrated: the 10%–90% credible intervals covered the simulated parameters 72% of the time. FPRs (false positive rates) were characterized by fixing *β*_2_ = *β*_3_ = 1 (no link between trait variation and molecular evolution) and simulating 200 times under two scenarios. In the first scenario (Table 1, col. 2) elements were generated under an exclusively neutral process, in the second scenario (Table 1, col. 5) completely conserved elements were generated. In the latter conserved scenario *r* = 0.2 was used, the expected value of *r*_2_ under our prior. The FPR dropped below 1% in the neutral scenario when a BF of 2 or more was used as a cut-off; under the conserved scenario the corresponding cut-off was also BF *≥* 2.

**Figure 1:**
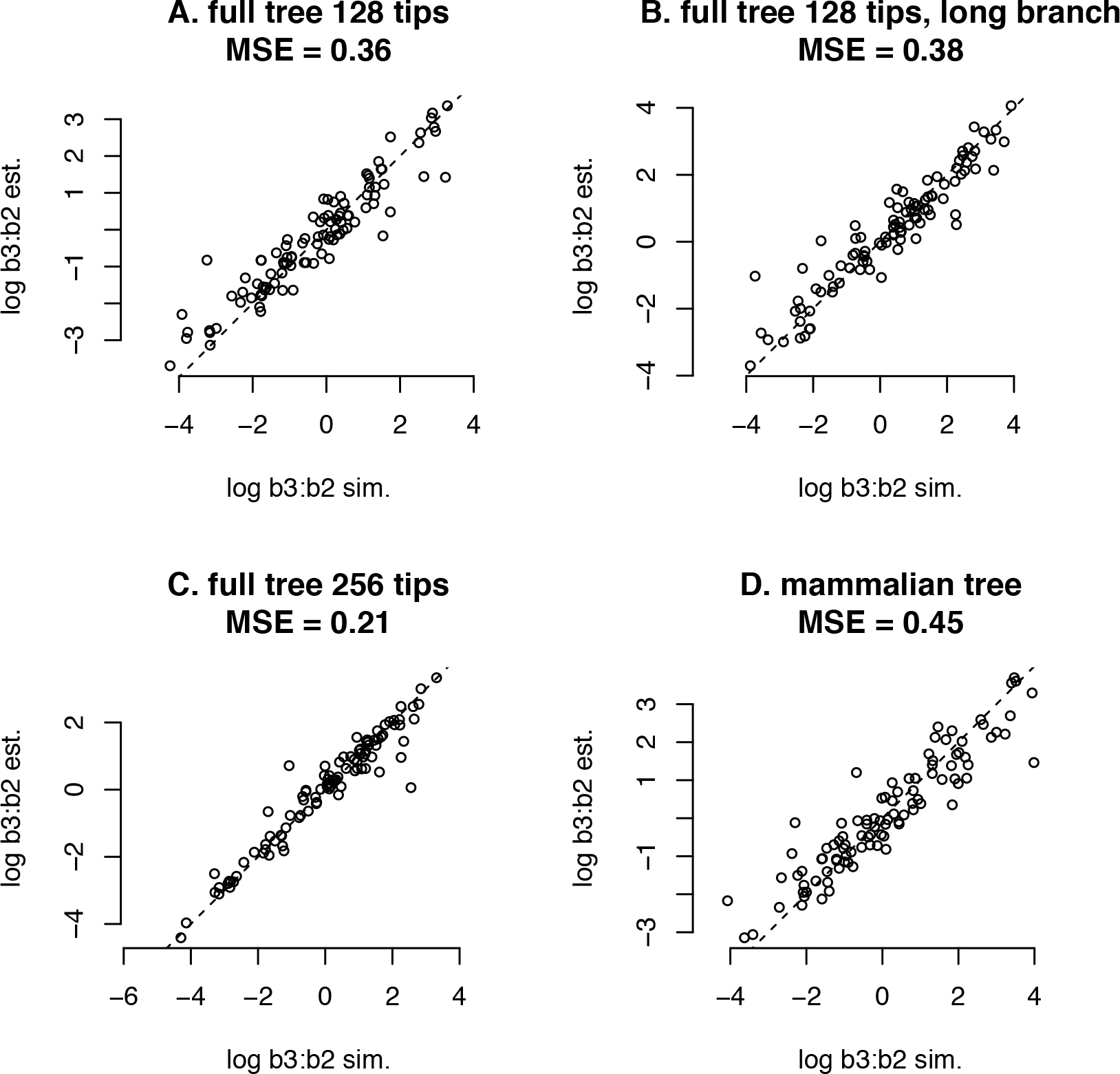
Simulated versus recovered (median) log *β*_3_ : *β*_2_ under model using 80 bp alignments. *A*. fully bifurcating ultrametric tree with 128 tips and all branch lengths set to 0.1; *B*. branch lengths are doubled to 0.2; *C*. tip count is doubled with respect to *A*, but branch lengths are reduced to 0.09 to keep root to tip distance similar; *D*. branch lengths and topology as per mammalian tree (see S1 Text).

**Table 1:** FPR when choosing the full model using the BF in favour as a cut-off. FPRs were estimated using 200 simulations for each of six scenarios. The tree variants were: 128 tips, branch length 0.1; 128 tips, branch length 0.2; 256 tips, branch length 0.09. The alignment variants used rate multiplier 1 (neutral) and 0.2 (conserved).

| BF cut-off | neutral<br>128 tips | neutral<br>128 tips<br>long branch | neutral<br>256 tips | conserved<br>128 tips | conserved<br>128 tips<br>long branch | conserved<br>256 tips |
| --- | --- | --- | --- | --- | --- | --- |
| 1 | 100.0% | 99.5% | 100.0% | 100.0% | 97.5% | 98.5% |
| 2 | 0.0% | 0.0% | 0.5% | 0.0% | 0.0% | 1.0% |
| 3 | 0.0% | 0.0% | 0.0% | 0.0% | 0.0% | 0.0% |

We took the bifurcating tree described above and doubled the branch lengths to 0.2 and then performed simulations analogous to those described above. We found we were no better able to recover log *β*_3_ : *β*_2_, now seeing an MSE of 0.38 (Figure 1B). The model remained reasonably calibrated as the 10%–90% credible intervals covered the simulated parameters 74% of the time. FPRs dropped to less than 1% by using BF cut-offs *≥* 2 in both scenarios (Table 1, cols. 3 and 6).

In the last of our idealized scenarios we doubled the number of tips on the tree to 256 while reducing the branch lengths to 0.09. In this scenario, recovery of log *β*_3_ : *β*_2_ improved, having an MSE of 0.21 (Figure 1C). The 10%–90% credible intervals covered the simulated parameters 85% of the time and BF cut-offs of 3 or more were sufficient to reduce the FPR to less than 1% (Table 1, cols. 4 and 7).

#### Simulations on a mammalian tree

Ultimately only performance on real topologies and with real traits matters. In a manner similar to the above simulations, we took our mammalian tree, having 61 tips, a variety of branch lengths, and an unbalanced topology, and simulated from our prior distributions as before. Recovery of log *β*_3_ : *β*_2_ worked less well, with an MSE of 0.45 (Figure 1D), though the model characterized its uncertainty appropriately as the 10%–90% credible intervals covered the simulated parameters 79% of the time.

When testing FPR, we considered scenarios relevant to our size and lifespan results (below). Therefore, we fixed the values of the trait to the real LLL trait values of Kowalczyk et al. [15] i.e. we simulated only conservation states and alignments. Three scenarios were devised: one in which all elements evolved neutrally, one in which elements were conserved at the expected level of *r* = 0.2, and one in which the elements were conserved at *r* = 0.5, which put them above the 99th percentile according to our prior. When simulating short elements (50 bp) against the LLL trait we needed BF cut-offs of 7 (neutral), 3 (conserved), and 8 (barely conserved) in order to reduce the FPR below 1% (Table 2, cols. 2–4); when considering longer elements (180 bp) the relevant BF thresholds were 4, 4, and 7 (Table 2, cols. 4–7). We remark that the barely conserved scenario (*r* = 0.5) presents a challenging set of parameters for the model, which does not mix well when *r*_2_ and *r*_3_ are conflated.

**Table 2:** FPR on the mammalian tree (see S1 Text) when choosing the full model using the BF in favour as a cut-off. FPRs were estimated using 200 simulations for each of six scenarios. The rate multiplier variants used to generate the alignments were: 1 (neutral); 0.2 (conserved); 0.5 (barely conserved). The alignment length variants were 50 bp and 180 bp. The trait was not simulated but fixed to the LLL values (see Figure 2 or S2 Tabular data).

| BF cut-off | neutral<br>50 bp | conserved<br>50 bp | barely<br>conserved<br>50 bp | neutral<br>180 bp | conserved<br>180 bp | barely<br>conserved<br>180 bp |
| --- | --- | --- | --- | --- | --- | --- |
| 1 | 100.0% | 100.0% | 100.0% | 100.0% | 100.0% | 100.0% |
| 2 | 19.5% | 1.5% | 65.0% | 1.5% | 2.5% | 37.5% |
| 3 | 2.0% | 0.0% | 10.5% | 0.5% | 1.5% | 5.5% |
| 4 | 1.5% | 0.0% | 2.5% | 0.0% | 0.0% | 2.0% |
| 5 | 1.0% | 0.0% | 0.5% | 0.0% | 0.0% | 1.0% |
| 6 | 0.5% | 0.0% | 0.5% | 0.0% | 0.0% | 0.5% |
| 7 | 0.0% | 0.0% | 0.5% | 0.0% | 0.0% | 0.0% |
| 8 | 0.0% | 0.0% | 0.0% | 0.0% | 0.0% | 0.0% |

### 3.2 Size and lifespan of mammals

Rather than pre-filtering CNE alignments based on heuristics, we instead ran PhyloAcc-C on all alignments with the LLL trait as input and then considered those where the model fit well in a reasonable 10,000 iterations, as assessed via a Gelman and Rubin [27] convergence diagnostic of *<* 1.01 across 3 chains. This resulted in summaries for 136,859 elements.

We ranked the elements by the BF (Bayes factor) in favour of the full model, where the rate of trait evolution is allowed to co-vary with the rate of molecular evolution, versus the null model, where the rate of trait evolution is constant across the phylogeny. We found 30 elements (0.02% of total) where the full model was ‘overwhelmingly’ supported (BF *≥* 100) with respect to the LLL trait and 1,109 (0.81% of total) where the full model was ‘very strongly’ supported (BF *≥* 30). We note that a BF *≥* 30 generally corresponded to effect sizes of magnitude 2 or more on the log scale i.e. to a ratio of about 7*×* or more.

The result of running PhyloAcc-C on the element with the highest BF is shown in Figure 2. The ancestral reconstruction of the LLL trait with respect to this element is shown in Figure 2 of S1 Text.

**Figure 2:**
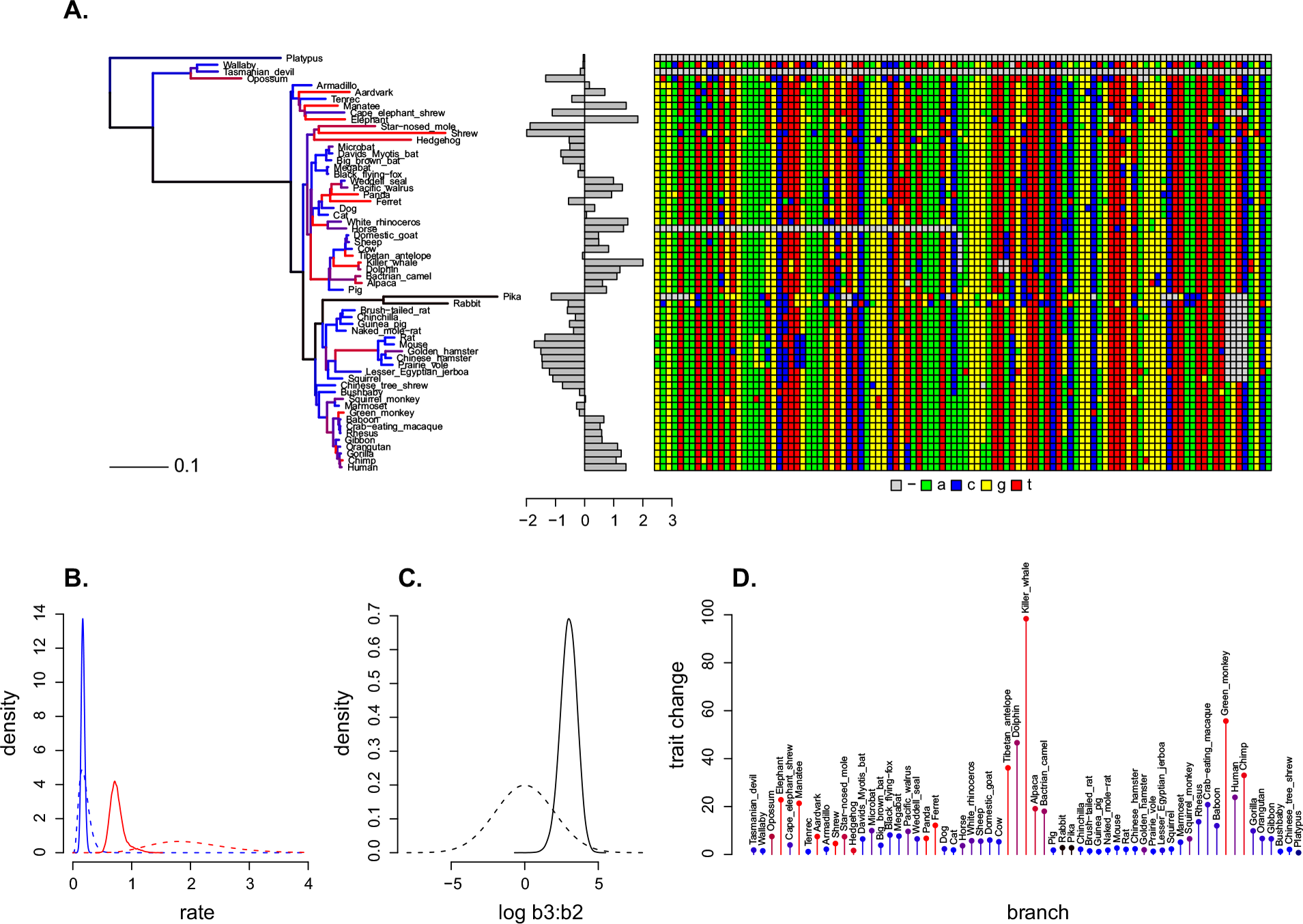
PhyloAcc-C fit to the LLL loci with the highest BF in favour of the full model (VCE277691). *A*. the mammalian phylogeny (input data, see S1 Text) is scaled according to the posterior distribution of rate multipliers **r** and coloured by the posterior distribution of conservation state **z** (black=neutral, blue=conserved, red=accelerated). Next to the tree the LLL trait and CNE alignment (both are also input data) are shown. The corresponding posterior distribution of the trait (i.e. an ancestral reconstruction) is shown in Figure 2 of S1 Text. *B*. the prior (dashed) and posterior (solid) distribution of the rate multipliers *r*_2_ (blue, conserved) and *r*_3_ (red, accelerated). *C*. the prior (dashed) and posterior (solid) distribution of log *β*_3_ : *β*_2_. In this case the posterior distribution suggests a positive value so that faster nucleotide evolution is associated with faster trait evolution, but see S1 Text for VCE351367 where the opposite is true. *D*. posterior distribution of trait change from tip to immediate ancestor, normalized by branch length and coloured by posterior conservation state. Again note that an accelerated conservation state (red) is associated with bigger trait moves and a conserved conservation state (blue) is associated with smaller ones.

To determine if there were biologically interesting patterns that could be systematically detected based on CNE location, we submitted the 1,109 loci of interest as genomic foreground to a GREAT analysis [28]; the GREAT tool ‘assigns biological meaning to a set of non-coding genomic regions by analyzing the annotations of the nearby genes’ and is a more principled alternative to an analysis based solely on the gene nearest a given CNE. The full set of CNEs (not the whole genome) were used as genomic background for the analysis.

The GREAT analysis suggested no genes were associated with the 1,109 LLL loci of interest, although several GO (gene ontology) biological processes were. These can be summarized as: blood vessel endothelial cell proliferation involved in sprouting angiogenesis; positive regulation of branching involved in lung morphogenesis; regulation of muscle tissue development, differentiation, and proliferation, esp. in the heart; regulation of alkaline phosphatase activity; astrocyte development; organ induction; endocrine pancreas development; trachea formation.

In addition to performing an analysis using GREAT, we also examined the 1 Mbp regions surrounding the top 25 loci associated with the LLL trait using the UCSC [10] and ENSEMBL [29] genome browsers, noting known functions or associations of nearby genes. To do this we made use of the GeneCards tool [30], and linked databases, such as the GWAS Catalog [31]. We found that 12 loci were near genes associated with height, weight, or limb length in some way, mainly via GWAS. Seven loci were associated with cancer genes, seven with the brain or nervous system, six with the skeleton, four with sperm, and one with longevity. Three regions had little to no annotation available whereas four loci were associated with p53, cell fate, or telomere length.

Overall, LLL loci with BF *≥* 30 exhibited effects in both directions, with log *β*_3_ : *β*_2_ being both positive and negative (Figure 3). The results of our note taking approach, and summaries of the PhyloAcc-C runs on the 136,859 LLL loci are recorded in S2 Tabular data. Full output of the GREAT analysis is reported in S3 Output of GREAT analysis on LLL candidate loci.

**Figure 3:**
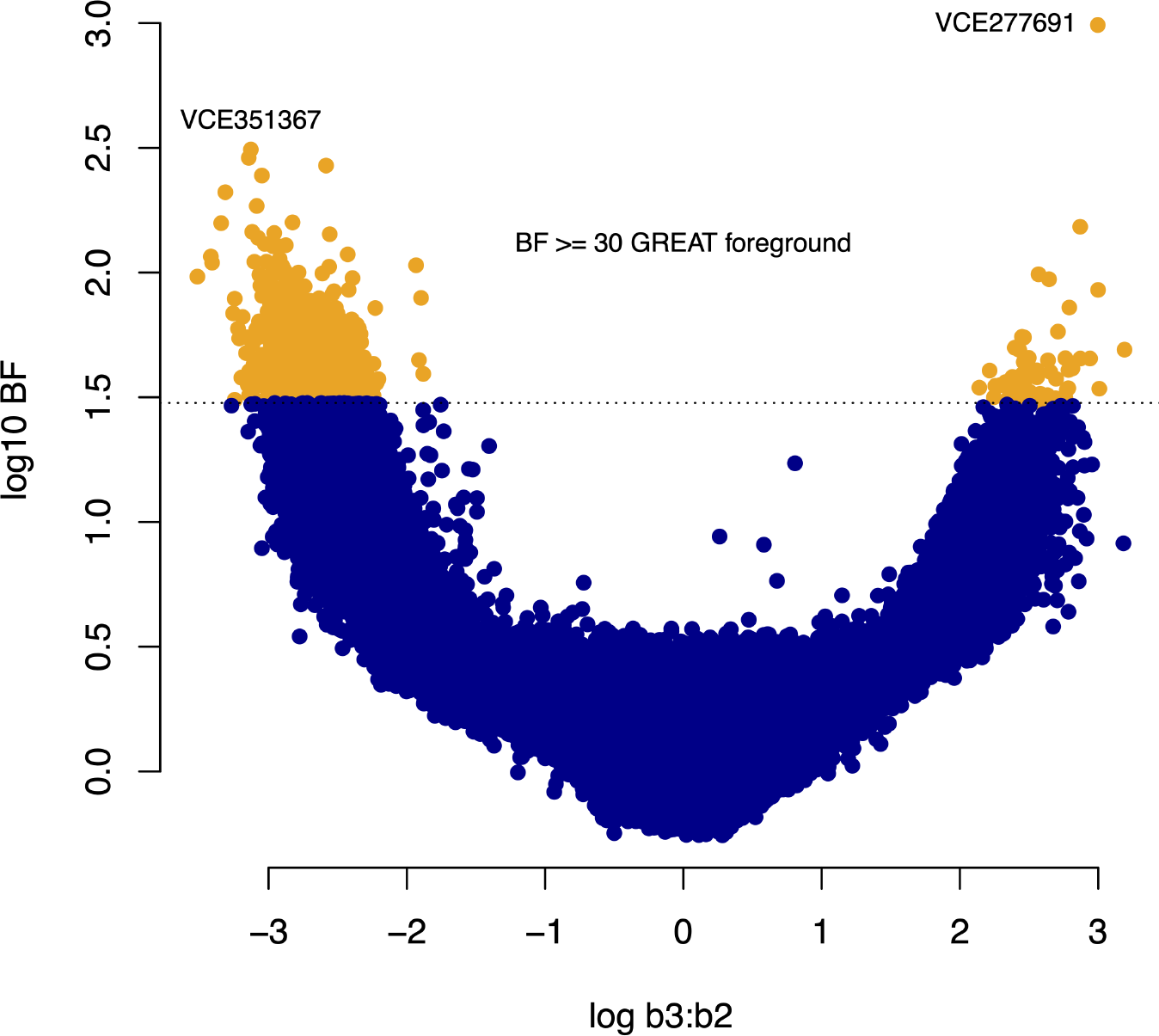
BF versus estimated (median) log *β*_3_ : *β*_2_ for 136,859 mammalian LLL loci. Orange loci are those having BF *≥* 30 and that were submitted as GREAT foreground during analysis. The two loci with the highest BF in favour of the full model are labelled. Note VCE277691 (see Figure 2) and VCE351367 (see S1 Text) have effects with opposite signs.

## 4 Discussion

We present a statistical method, PhyloAcc-C, for relating the rate of nucleotide evolution to the rate of evolution of a continuous trait. The model is phylogenetically framed and operates under the common assumption that nucleotide evolution follows a site-independent, continuous-time discrete-state Markov process, and that continuous traits evolve under Brownian motion, though in our case with potentially different rates on different parts of the tree. Latent rate categories are also assigned using a Markov process, which all together allows the rates of molecular and phenotypic evolution to vary in an automatic way across branches.

A notable feature of the model is its ability to associate the evolution of continuous traits and non-coding DNA using a more statistically integrated approach than that taken by Yusuf et al. [14] or Kowalczyk et al. [15]. Indeed, the general idea of linking genotypic rate multipliers (i.e. evolution relative to a known background tree) and phenotypic rate multipliers (i.e. variance parameters) seems natural, and could be used in other frameworks. For example, the PhyloAcc family of models (https://phyloacc.github.io/) allocates rates in a way which was designed with efficient processing of CNEs in mind, yet genotypic and phenotypic rates of evolution can also be linked under more complex models, such as relaxed clocks [32] or local clocks [33, 34], via linear or logistic functions. The efficiency versus accuracy tradeoffs of different rate assignment strategies will not be clear without further research.

Simulations show that the model can perform acceptably on both ideal trees and a mammalian tree related to a large set of CNE alignments. This is encouraging, but we suggest users of the method check model performance using their tree, and the sequence lengths and rate multipliers they expect to see. The R software package and instructions accompanying this paper make the simulation process relatively straightforward. In addition, we emphasize that whereas larger trees might provide more data points, there is a judgment call to be made over how large a tree can be plausibly described by three rate categories.

As an illustrative example, we applied PhyloAcc-C to CNEs and longevity data from mammals. From a biomedical perspective, longevity is an important trait, with a long history of study in a diversity of organisms, from worms [35] to humans [36], and therefore there is at least some possibility of assessing the plausibility of candidate loci using published evidence and annotations. Furthermore, lifespan and body-size are also relevant to longstanding conundrums and current ecological debates, including theories of life-history tradeoffs [37, 38], and Peto’s paradox [39], which asks how large and long-lived animals mitigate cancer risk in the face of the many cell divisions that occur during their lifetime. This means the trait is also familiar and of interest to a broad readership. For these reasons, we describe three recent papers studying lifespan that also use broad genomic data, and help put our analysis and methodology into context.

Kowalczyk et al. [15] studied the LLL trait (as previously mentioned, we reuse their trait data) but in the context of protein evolution, with an explicit focus on genes that were interpreted as being under increased purifying selection in long-lived large-bodied species, where LLL was treated as the derived state. Kowalczyk et al. [15] highlighted processes related to the cell cycle, DNA repair, cell death, immunity, and IGF1 expression pathways. Each of these processes were then plausibly linked to lifespan via their analysis. The authors also discuss telomere maintenance and p53, also plausibly linked to aging and cancer control. There is notable overlap between our LLL results and theirs: we also found associations to p53, telomere maintenance, and cell fate within 1 Mbp of our top 25 loci of interest. Our top 25 loci also have links to cancer and height or body size, though these prevalent diseases and biomarkers are of course heavily studied and consequently commonly annotated, and so we cannot know whether their appearance is simply due to their frequency.

Tejada-Martinez et al. [40] also focused on protein evolution in their study of lifespan and body mass in primates, although they then linked their findings to enhancer evolution. Their approach was to perform phylogenetic regressions, relating dN/dS to maximum lifespan and body mass for around 10,000 genes. In contrast to Kowalczyk et al. [15], Tejada-Martinez et al. [40] focus on positive (directional) selection on protein-coding genes rather than conservation. The authors identified 276 candidate genes whose rate of adaptive evolution positively correlated with maximum life span in a phylogenetic context. The authors focused their discussion on the enrichment of diverse processes including immunity, inflammation, cellular aging, organismal development (height, BMI), neurodevelopment, and brain function. These processes are all represented in our results in one form or another. None of the genes mentioned in the body of their manuscript occur in our notes on our top 25 loci of interest except for p53, the well known tumor suppressor, which is downregulated by HDAC3, a gene close to the CNE ranked as most-interesting overall in our LLL analysis (Figure 2).

A third study by Treaster et al. [41] takes yet a different approach to understanding longevity. By focusing on 23 species of rockfish that are both closely related and feature a wide range of lifespans (11 to more than 200 years), the authors aimed to identify longevity related protein coding genes while minimizing false positives due to (other) convergent evolution. A key part of the analysis pipeline was the detection of rate shifts using the TRACCER tool [7]. Unlike Kolora et al. [42], and the other studies we mention, Treaster et al. [41] treated longevity as a binary trait and argued against the need to correct for body size. The authors found the ancestral rockfish state to be long-lived, and linked positive selection to glycogen biosynthesis and flavonoid metabolism via GO analysis. The top genes identified in their study do not feature in our notes on our top 25 LLL loci though, as all of us do, the authors find a relationship between their loci of interest and p53, in this case via PLA2R1. We note Treaster et al. [41] and Kowalczyk et al. [15] emphasize insulin signaling pathways, though apparently the particular pathways are under increased constraint in mammals (gene IGF1) but accelerated in Rockfish (gene INSR).

The above studies either focus exclusively on protein coding genes [15, 41] or examine non-coding sequences only insofar as they are identified as byproduct of an analysis with genes as the starting point and main consideration [40]. One distinguishing factor of PhyloAcc-C when compared to these approaches is that its focus is on identifying relevant non-coding sequences. It is possible then that PhyloAcc-C will sometimes identify processes that would otherwise be missed. Indeed, Treaster et al. [41] specifically mention that an attempt was made to analyze the CNE data captured as part of their study, but that their approach was underpowered when working with short conserved sequences. This suggests PhyloAcc-C might be used in a complementary manner to existing methodologies, potentially extracting further insight from a given sequencing data set. Bearing this in mind, it is interesting that our analysis appeared to highlight alternative biological themes that are not present in the results of the above studies, but that do seem plausibly related to longevity and body size. These themes are a prevalence of associations with skeletal genes and genes relating to exploratory process.

When examining our top 25 LLL loci we noticed several genes related to bone strength or bone development including Fibrillin 2 (FBN2). We note that FBN2 is specifically associated with contractural arachnodactyly, i.e. a particularly tall long-limbed phenotype, with long slender fingers and toes. Such non-lethal but body size-related phenotypic differences do seem to be the kind of effects that one would a priori imagine to be associated with true LLL loci. In the case of exploratory processes, we note our GO analysis identified the processes ‘blood vessel endothelial cell proliferation involved in sprouting angiogenesis’ and ‘positive regulation of branching involved in lung morphogenesis’. These sort of developmental processes are exactly those thought to enhance evolvability [43, 44]. The basic reasoning is that while core functions are conserved across metazoa, the evolutionary flexibility of anatomical traits, such as limb shape or size, is derived from the fact that many of their constituent components are decoupled from a few fixed genetically coded features. For example, the limb is a co-ordinated collection of bone, muscle, nerves, and vasculature, but the genetic orchestration of limb development is largely achieved through cartilaginous condensations, which then select feasible arrangements of the aforementioned components. However, what works for the body during development can work against it in the case of cancer, and the importance of blood supply to tumour growth and metastasis means that as of 2018 at least 14 endothelial angiogenesis inhibitors were being used to treat cancer in the USA [45].

In conclusion then, we have introduced a method that can be used to study the co-evolution of continuous traits and non-coding DNA. The method is available as an R package and users are free to modify it as they wish under the GPL. Applying the method highlighted interesting candidate LLL loci, including those related to exploratory processes, skeletal development, as well as more ‘typical’ lifespan related themes that have also been identified in other recent bioinformatics studies.

We have given some thought to future work. Longevity and size are clearly complex traits that are both correlated with each other, and also with other traits, e.g. sociality [46]. Moreover, is not unreasonable to think that thousands of enhancers and (at least) hundreds of genes are systematically involved in the evolution of lifespan and body size. PhyloAcc-C has the weakness that it cannot tease apart the relative contribution of different loci to traits of interest. One future direction then would be to focus on methods for finding clusters of loci that collectively, but not always simultaneously, contribute to the variation of a trait. Another area for future work is the incorporation of more flexible null models. One way this should be attempted is by using a more realistic method to assign nucleotide rate multipliers to branches, such as relaxed, correlated, or random local clocks, or their more recent derivatives e.g. [47]. A second improvement would be to make use of alternative models of trait evolution such as those used by Uyeda and Harmon [48]. A combination of more flexible rate assignment and alternative models of trait evolution would lead to a more plausible null model overall, giving greater confidence that a high BF indicates an interesting locus.

In the mean time, we see our method, and the other PhyloG2P methods we have discussed, as first steps towards powerful tools to advance the PhyloG2P programme. Such methods will ultimately increase both our understanding of natural history and also allow us to use data from diverse species to shine a spotlight on parts of our own genome that are important for biodiversity and human health. Smith et al. [1] put it well: ‘Phylogenetics is the new genetics’.

## Supporting information

S2. Tabular data.

S3. Output of GREAT analysis on LLL candidate loci.

## 5 Data and software availability

The alignments and tree used in this study are those previously used and made available by Hu et al. [12]. The trait data used in this study is collated by Kowalczyk et al. [15]. The PhyloAcc-C software can be found at https://github.com/phyloacc.

## 6 Supplementary files

S1. Text. Additional text and figures. S2. Tabular data. S3. Output of GREAT analysis on LLL candidate loci.

## Acknowledgements

We thank members of the Edwards, Liu, and Sackton groups for helpful discussions during the course of this research. The computations in this paper were run on the FASRC Cannon cluster supported by the FAS Division of Science Research Computing Group at Harvard University.

## References

[1] Smith SD, Pennell MW, Dunn CW, Edwards SV. Phylogenetics is the new genetics (for most of biodiversity). Trends in Ecology & Evolution. 2020;35(5):415–25.

[2] Hiller M, Schaar BT, Indjeian VB, Kingsley DM, Hagey LR, Bejerano G. A “forward genomics” approach links genotype to phenotype using independent phenotypic losses among related species. Cell Reports. 2012;2(4):817–23.

[3] Marcovitz A, Jia R, Bejerano G. “Reverse genomics” predicts function of human conserved noncoding elements. Molecular Biology and Evolution. 2016;33(5):1358–69.

[4] Prudent X, Parra G, Schwede P, Roscito JG, Hiller M. Controlling for phylogenetic relatedness and evolutionary rates improves the discovery of associations between species’ phenotypic and genomic differences. Molecular Biology and Evolution. 2016;33(8):2135–50.

[5] Langer BE, Roscito JG, Hiller M. REforge associates transcription factor binding site divergence in regulatory elements with phenotypic differences between species. Molecular Biology and Evolution. 2018;35(12):3027–40.

[6] Partha R, Kowalczyk A, Clark NL, Chikina M. Robust method for detecting convergent shifts in evolutionary rates. Molecular Biology and Evolution. 2019;36(8):1817–30.

[7] Treaster S, Daane JM, Harris MP. Refining convergent rate analysis with topology in mammalian longevity and marine transitions. Molecular Biology and Evolution. 2021;38(11):5190–203.

[8] Hardison RC. Conserved noncoding sequences are reliable guides to regulatory elements. Trends in Genetics. 2000;16(9):369–72.

[9] Siepel A, Pollard KS, Haussler D. New methods for detecting lineage-specific selection. In: Research in Computational Molecular Biology: 10th Annual International Conference, RECOMB 2006, Venice, Italy, April 2-5, 2006. Proceedings 10. Springer; 2006. p. 190–205.

[10] Rosenbloom KR, Armstrong J, Barber GP, Casper J, Clawson H, Diekhans M, et al. The UCSC genome browser database: 2015 update. Nucleic Acids Research. 2015;43(D1):D670–81.

[11] Booker BM, Friedrich T, Mason MK, VanderMeer JE, Zhao J, Eckalbar WL, et al. Bat accelerated regions identify a bat forelimb specific enhancer in the HoxD locus. PLoS Genetics. 2016;12(3):e1005738.

[12] Hu Z, Sackton TB, Edwards SV, Liu JS. Bayesian detection of convergent rate changes of conserved noncoding elements on phylogenetic trees. Molecular Biology and Evolution. 2019;36(5):1086–100.

[13] Lartillot N, Poujol R. A phylogenetic model for investigating correlated evolution of substitution rates and continuous phenotypic characters. Molecular Biology and Evolution. 2011;28(1):729–44.

[14] Yusuf L, Heatley MC, Palmer JP, Barton HJ, Cooney CR, Gossmann TI. Noncoding regions underpin avian bill shape diversification at macroevolutionary scales. Genome Research. 2020;30(4):553–65.

[15] Kowalczyk A, Partha R, Clark NL, Chikina M. Pan-mammalian analysis of molecular constraints underlying extended lifespan. Elife. 2020;9:e51089.

[16] Pollard KS, Hubisz MJ, Rosenbloom KR, Siepel A. Detection of nonneutral substitution rates on mammalian phylogenies. Genome Research. 2010;20(1):110–21.

[17] Liu JS. The collapsed Gibbs sampler in Bayesian computations with applications to a gene regulation problem. Journal of the American Statistical Association. 1994;89(427):958–66.

[18] Liu J. Monte Carlo strategies in scientific computing. Statistics, Springer-Verlag, New York. 2001.

[19] Felsenstein J. Evolutionary trees from DNA sequences: a maximum likelihood approach. Journal of Molecular Evolution. 1981;17:368–76.

[20] Wagenmakers EJ, Lodewyckx T, Kuriyal H, Grasman R. Bayesian hypothesis testing for psychologists: A tutorial on the Savage–Dickey method. Cognitive Psychology. 2010;60(3):158–89.

[21] Blanchette M, Kent WJ, Riemer C, Elnitski L, Smit AF, Roskin KM, et al. Aligning multiple genomic sequences with the threaded blockset aligner. Genome Research. 2004;14(4):708–15.

[22] Murphy WJ, Pevzner PA, O’Brien SJ. Mammalian phylogenomics comes of age. Trends in Genetics. 2004;20(12):631–9.

[23] R Core Team. R: A Language and Environment for Statistical Computing. Vienna, Austria; 2021. Available from: https://www.R-project.org/.

[24] Eddelbuettel D, Francois R. Rcpp: Seamless R and C++ integration. Journal of Statistical Software. 2011;40:1–18.

[25] Eddelbuettel D, Sanderson C. RcppArmadillo: Accelerating R with high-performance C++ linear algebra. Computational Statistics & Data Analysis. 2014;71:1054–63.

[26] Paradis E, Claude J, Strimmer K. APE: analyses of phylogenetics and evolution in R language. Bioinformatics. 2004;20(2):289–90.

[27] Gelman A, Rubin DB. Inference from iterative simulation using multiple sequences. Statistical Science. 1992:457–72.

[28] McLean CY, Bristor D, Hiller M, Clarke SL, Schaar BT, Lowe CB, et al. GREAT improves functional interpretation of cis-regulatory regions. Nature Biotechnology. 2010;28(5):495–501.

[29] Flicek P, Amode MR, Barrell D, Beal K, Brent S, Carvalho-Silva D, et al. Ensembl 2012. Nucleic Acids Research. 2012;40(D1):D84–90.

[30] Stelzer G, Rosen N, Plaschkes I, Zimmerman S, Twik M, Fishilevich S, et al. The GeneCards suite: from gene data mining to disease genome sequence analyses. Current Protocols in Bioinformatics. 2016;54(1):1–30.

[31] Sollis E, Mosaku A, Abid A, Buniello A, Cerezo M, Gil L, et al. The NHGRI-EBI GWAS Catalog: knowledgebase and deposition resource. Nucleic Acids Research. 2023;51(D1):D977–85.

[32] Drummond AJ, Ho SYW, Phillips MJ, Rambaut A. Relaxed phylogenetics and dating with confidence. PLoS Biology. 2006;4(5):e88.

[33] Thorne JL, Kishino H, Painter IS. Estimating the rate of evolution of the rate of molecular evolution. Molecular Biology and Evolution. 1998;15(12):1647–57.

[34] Drummond AJ, Suchard MA. Bayesian random local clocks, or one rate to rule them all. BMC Biology. 2010;8(1):1–12.

[35] Dorman JB, Albinder B, Shroyer T, Kenyon C. The age-1 and daf-2 genes function in a common pathway to control the lifespan of Caenorhabditis elegans. Genetics. 1995;141(4):1399–406.

[36] Deelen J, Evans DS, Arking DE, Tesi N, Nygaard M, Liu X, et al. A meta-analysis of genome-wide association studies identifies multiple longevity genes. Nature Communications. 2019;10(1):3669.

[37] Maklakov AA, Immler S. The expensive germline and the evolution of ageing. Current Biology. 2016;26(13):R577–86.

[38] Muntané G, Farré X, Rodríguez JA, Pegueroles C, Hughes DA, de Magalhaes JP, et al. Biological processes modulating longevity across primates: a phylogenetic genomephenome analysis. Molecular Biology and Evolution. 2018;35(8):1990–2004.

[39] Tollis M, Boddy AM, Maley CC. Peto’s Paradox: how has evolution solved the problem of cancer prevention? BMC Biology. 2017;15:1–5.

[40] Tejada-Martinez D, Avelar RA, Lopes I, Zhang B, Novoa G, De Magalhaes JP, et al. Positive selection and enhancer evolution shaped lifespan and body mass in great apes. Molecular Biology and Evolution. 2022;39(2):msab369.

[41] Treaster S, Deelen J, Daane JM, Murabito J, Karasik D, Harris MP. Convergent genomics of longevity in rockfishes highlights the genetics of human life span variation. Science Advances. 2023;9(2):eadd2743.

[42] Kolora SRR, Owens GL, Vazquez JM, Stubbs A, Chatla K, Jainese C, et al. Origins and evolution of extreme life span in Pacific Ocean rockfishes. Science. 2021;374(6569):842–7.

[43] Kirschner M, Gerhart J. Evolvability. Proceedings of the National Academy of Sciences. 1998;95(15):8420–7.

[44] Kirschner MW, Gerhart JC. The plausibility of life: Resolving Darwin’s dilemma. Yale University Press; 2005.

[45] NIH National Cancer Institute. Angiogenesis Inhibitors; 2018. [Online; accessed 1-August-2023]. https://www.cancer.gov/about-cancer/treatment/types/immunotherapy/angiogenesis-inhibitors-fact-sheet.

[46] Zhu P, Liu W, Zhang X, Li M, Liu G, Yu Y, et al. Correlated evolution of social organization and lifespan in mammals. Nature Communications. 2023;14(1):372.

[47] Fisher AA, Ji X, Nishimura A, Lemey P, Suchard MA. Shrinkage-based random local clocks with scalable inference. arXiv preprint arXiv:210507119. 2021.

[48] Uyeda JC, Harmon LJ. A novel Bayesian method for inferring and interpreting the dynamics of adaptive landscapes from phylogenetic comparative data. Systematic Biology. 2014;63(6):902–18.

