## Supplementary material for "A phylogenetic method linking nucleotide substitution rates to rates of continuous trait evolution": S1. Text. Additional text and figures.

### DRAFT: S1 Text

We used the following rate matrix and stationary distribution to model the background rate of nucleotide evolution:

$$\mathbf{Q} = \begin{pmatrix} -1.075162 & 0.186970 & 0.696268 & 0.191923 \\ 0.181082 & -0.873473 & 0.255492 & 0.436899 \\ 0.674340 & 0.255493 & -1.164645 & 0.234813 \\ 0.191924 & 0.451108 & 0.242449 & -0.885481 \end{pmatrix}, \pi = \begin{pmatrix} 0.246 \\ 0.254 \\ 0.254 \\ 0.246 \end{pmatrix}.$$

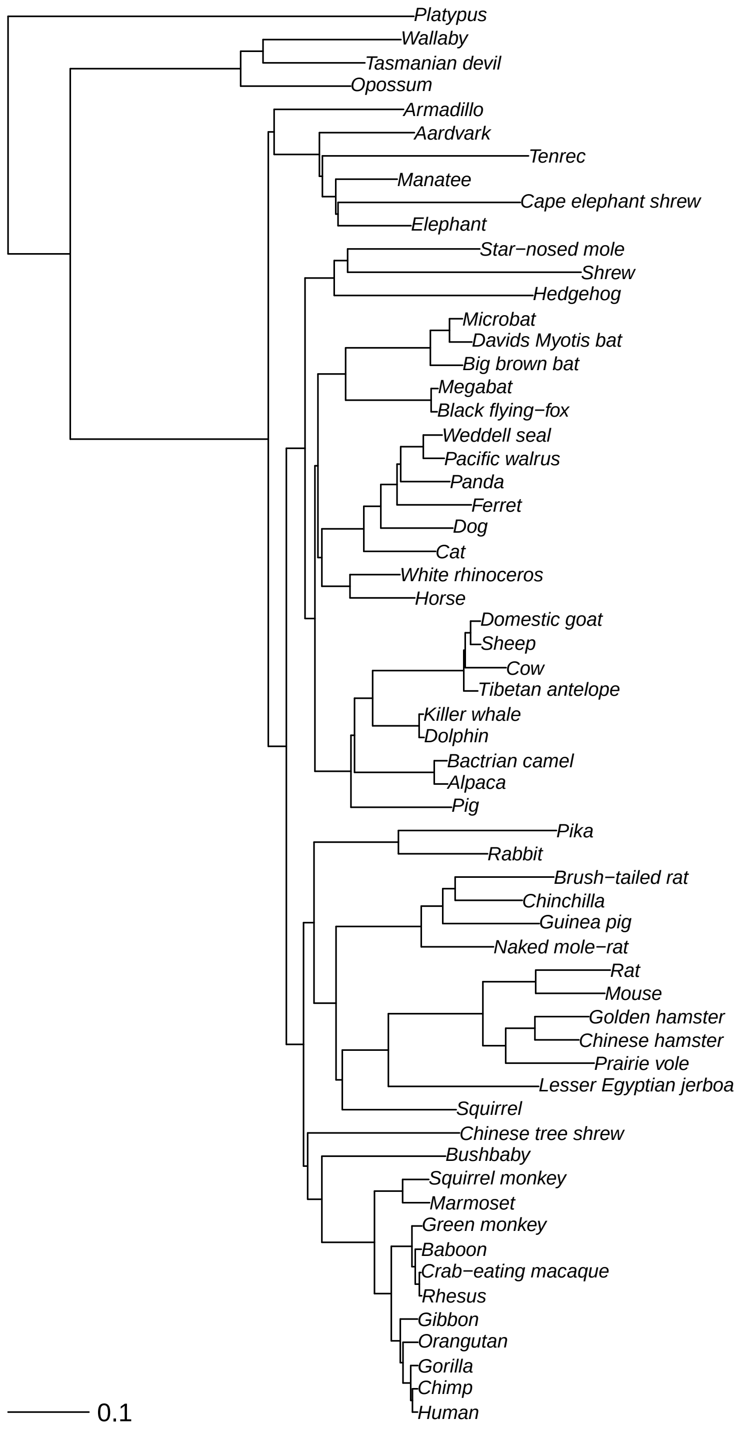


Figure 1: The tree used to model the background relationship between mammalian species. The tree was obtained from the PhyloAcc GitHub repository prepared by Hu et al. [1], available at: https://github.com/phyloacc/Hu-etal-2019-data/. The mammal phylogeny was originally prepared by Murphy et al. [2]. Note, we removed the Cape golden mole as it did not appear in the Kowalczyk et al. [3] dataset. The mapping from tips to the LLL trait data of [3] is show in S2 Tabular data.

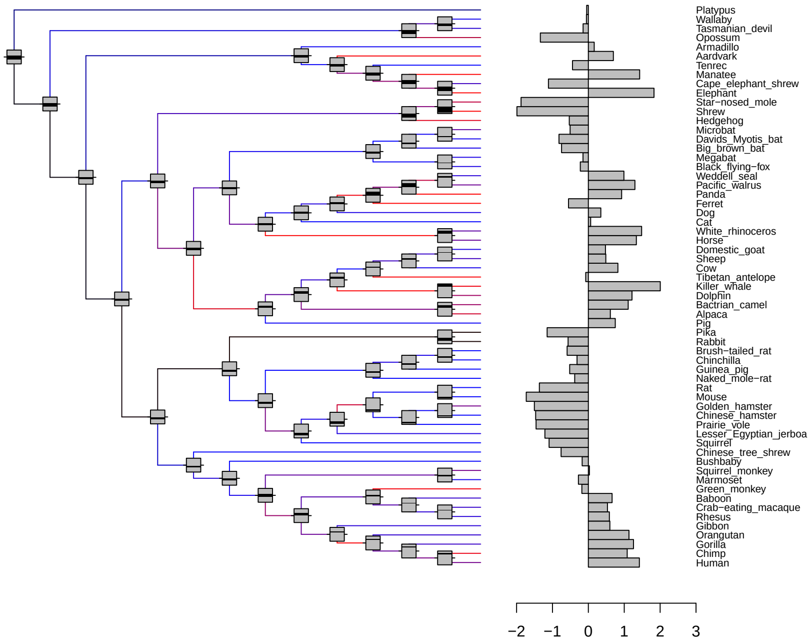


Figure 2: Ancestral reconstruction of the LLL trait for locus VCE277691. Black bars inside grey boxes at nodes show 10%–90% posterior reconstruction of trait. Grey boxes at nodes cover range from minimum of all 10% intervals to maximum of all 90% intervals.

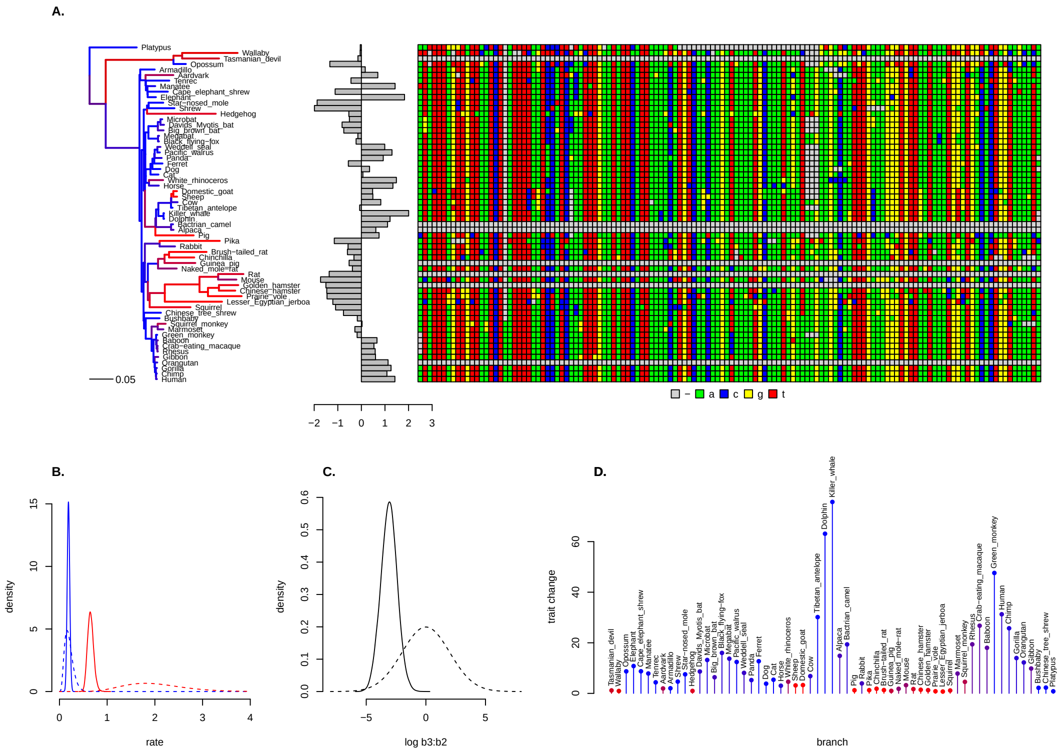


Figure 3: PhyloAcc-C fit to the LLL loci with the second highest BF in favour of the full model (VCE351367). *A.* the mammalian phylogeny (input data) is scaled according to the posterior distribution of rate multipliers $\mathbf{r}$ and coloured by the posterior distribution of conservation state $\mathbf{z}$ (black=neutral, blue=conserved, red=accelerated). Next to the tree the LLL trait and CNE alignment (both are also input data) are shown. The corresponding posterior distribution of the trait (i.e. an ancestral reconstruction) is shown in Figure 4. *B.* the prior (dashed) and posterior (solid) distribution of the rate multipliers $r_2$ (blue, conserved) and $r_3$ (red, accelerated). *C.* the prior (dashed) and posterior (solid) distribution of $\log \beta_3 : \beta_2$. In this case the posterior distribution suggests a negative value so that faster nucleotide evolution is associated with slower trait evolution. *D.* posterior distribution of trait change from tip to immediate ancestor, normalized by branch length and coloured by posterior conservation state. Again note that an accelerated conservation state (red) is associated with smaller trait moves and a conserved conservation state (blue) is associated with larger ones.

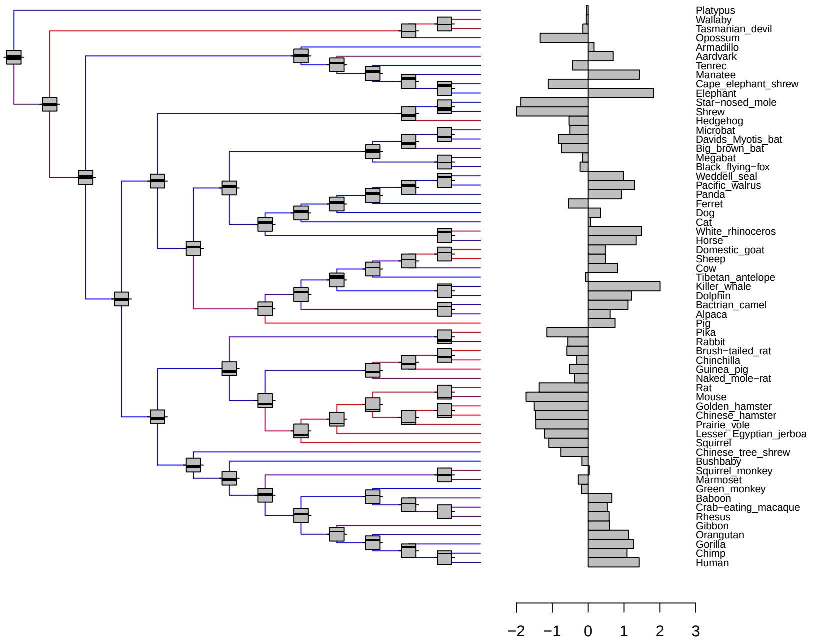


Figure 4: Ancestral reconstruction of the LLL trait for locus VCE351367. Black bars inside grey boxes at nodes show 10%–90% posterior reconstruction of trait. Grey boxes at nodes cover range from minimum of all 10% intervals to maximum of all 90% intervals.

### References


[1] Hu Z, Sackton TB, Edwards SV, Liu JS. Bayesian detection of convergent rate changes of conserved noncoding elements on phylogenetic trees. Molecular Biology and Evolution. 2019;36(5):1086-100.

[2] Murphy WJ, Pevzner PA, O'Brien SJ. Mammalian phylogenomics comes of age. Trends in Genetics. 2004;20(12):631-9.

[3] Kowalczyk A, Partha R, Clark NL, Chikina M. Pan-mammalian analysis of molecular constraints underlying extended lifespan. Elife. 2020;9:e51089.
