## Supplementary material for "A phylogenetic method linking nucleotide substitution rates to rates of continuous trait evolution": S3. Output of GREAT analysis on LLL candidate loci.

| Ontology | # Term Name | Term ID | Hyper Rank | Hyper Raw P-Value | Hyper Bonferroni P-Value | Hyper FDR Q-Val | Hyper Fold Enrichment | Hyper Expected | Hyper Foreground Region Hits | Hyper Total Regions | Hyper Region Set Coverage | Hyper Term Region Coverage | Hyper Foreground Gene Hits | Hyper Background Gene Hits | Total Genes Annotated |
|---|---|---|---|---|---|---|---|---|---|---|---|---|---|---|---|
| Ensembl Genes | No results meet your chosen criteria. | | | | | | | | | | | | | | |
| GO Biological Process | positive regulation of cardiocyte differentiation | GO:1905209 | 1 | 1.28778e-9 | 1.69459e-5 | 1.69459e-5 | 3.2619 | 11.0366 | 36 | 1362 | 3.25% | 2.64% | 9 | 22 | 25 |
| | positive regulation of cardiac muscle tissue development | GO:0055025 | 2 | 1.77092e-8 | 2.33035e-4 | 1.16518e-4 | 2.5377 | 18.1269 | 46 | 2237 | 4.15% | 2.06% | 13 | 36 | 40 |
| | positive regulation of striated muscle tissue development | GO:0045844 | 3 | 2.39969e-8 | 3.15775e-4 | 1.05258e-4 | 2.3309 | 22.7377 | 53 | 2806 | 4.78% | 1.89% | 16 | 56 | 64 |
| | positive regulation of cardiac muscle cell differentiation | GO:2000727 | 4 | 4.85776e-8 | 6.39232e-4 | 1.59808e-4 | 3.2683 | 8.8730 | 29 | 1095 | 2.61% | 2.65% | 7 | 14 | 17 |
| | blood vessel endothelial cell proliferation involved in sprouting angiogenesis | GO:0002043 | 6 | 2.06984e-7 | 2.72370e-3 | 4.53949e-4 | 5.3811 | 2.7875 | 15 | 344 | 1.35% | 4.36% | 2 | 7 | 7 |
| | positive regulation of branching involved in lung morphogenesis | GO:0061047 | 7 | 2.60776e-7 | 3.43155e-3 | 4.90221e-4 | 5.7020 | 2.4553 | 14 | 303 | 1.26% | 4.62% | 3 | 4 | 4 |
| | regulation of cardiocyte differentiation | GO:1905207 | 9 | 4.02047e-7 | 5.29054e-3 | 5.87838e-4 | 2.4082 | 17.0249 | 41 | 2101 | 3.70% | 1.95% | 13 | 36 | 41 |
| | positive regulation of muscle tissue development | GO:1901863 | 11 | 9.21456e-7 | 1.21254e-2 | 1.10231e-3 | 2.0692 | 25.6143 | 53 | 3161 | 4.78% | 1.68% | 16 | 57 | 65 |
| | regulation of cell proliferation involved in outflow tract morphogenesis | GO:1901963 | 12 | 1.18731e-6 | 1.56238e-2 | 1.30198e-3 | 5.9474 | 2.0177 | 12 | 249 | 1.08% | 4.82% | 2 | 2 | 2 |
| | negative regulation of | GO:2000242 | 14 | 3.08082e-6 | 4.05406e-2 | 2.89575e-3 | 2.7744 | 9.7320 | 27 | 1201 | 2.43% | 2.25% | 8 | 31 | 54 |

| | | | | | | | | | | | | | | | |
|---|---|---|---|---|---|---|---|---|---|---|---|---|---|---|---|
| | reproductive process | | | | | | | | | | | | | | |
| | regulation of cardiac muscle cell differentiation | GO:2000725 | 15 | 3.99040e-6 | 5.25097e-2 | 3.50064e-3 | 2.4415 | 13.5162 | 33 | 1668 | 2.98% | 1.98% | 10 | 24 | 29 |
| | smooth muscle cell differentiation | GO:0051145 | 16 | 4.41017e-6 | 5.80334e-2 | 3.62709e-3 | 2.3922 | 14.2131 | 34 | 1754 | 3.07% | 1.94% | 12 | 30 | 32 |
| | tendon development | GO:0035989 | 17 | 5.50877e-6 | 7.24899e-2 | 4.26411e-3 | 5.6327 | 1.9529 | 11 | 241 | 0.99% | 4.56% | 2 | 4 | 5 |
| | regulation of alkaline phosphatase activity | GO:0010692 | 18 | 5.64689e-6 | 7.43074e-2 | 4.12819e-3 | 3.7001 | 4.5945 | 17 | 567 | 1.53% | 3.00% | 4 | 7 | 9 |
| | astrocyte development | GO:0014002 | 19 | 6.79312e-6 | 8.93907e-2 | 4.70477e-3 | 3.4872 | 5.1618 | 18 | 637 | 1.62% | 2.83% | 4 | 20 | 27 |
| | organ induction | GO:0001759 | 20 | 7.32160e-6 | 9.63449e-2 | 4.81725e-3 | 2.9171 | 7.8844 | 23 | 973 | 2.07% | 2.36% | 6 | 15 | 15 |
| | positive regulation of cardiac muscle tissue growth | GO:0055023 | 21 | 8.15244e-6 | 1.07278e-1 | 5.10847e-3 | 2.3948 | 13.3622 | 32 | 1649 | 2.89% | 1.94% | 8 | 27 | 30 |
| | cardiac muscle cell differentiation | GO:0055007 | 22 | 9.00607e-6 | 1.18511e-1 | 5.38686e-3 | 2.0357 | 22.1056 | 45 | 2728 | 4.06% | 1.65% | 18 | 64 | 77 |
| | endocrine pancreas development | GO:0031018 | 23 | 9.13124e-6 | 1.20158e-1 | 5.22426e-3 | 2.2176 | 16.6846 | 37 | 2059 | 3.34% | 1.80% | 12 | 32 | 38 |
| | trachea formation | GO:0060440 | 24 | 9.13554e-6 | 1.20215e-1 | 5.00894e-3 | 4.8714 | 2.4634 | 12 | 304 | 1.08% | 3.95% | 2 | 6 | 6 |
| GO Cellular Component | BBSome | GO:0034464 | 1 | 9.29158e-6 | 1.60558e-2 | 1.60558e-2 | 5.9331 | 1.6855 | 10 | 208 | 0.90% | 4.81% | 3 | 9 | 10 |
| | ciliary transition zone | GO:0035869 | 2 | 9.85734e-6 | 1.70335e-2 | 8.51674e-3 | 3.5438 | 4.7971 | 17 | 592 | 1.53% | 2.87% | 7 | 37 | 56 |
| GO Molecular Function | No results meet your chosen criteria. | | | | | | | | | | | | | | |
| Human Phenotype | Abdominal distention | HP:0003270 | 1 | 5.85501e-7 | 3.90705e-3 | 3.90705e-3 | 3.6501 | 5.7533 | 21 | 710 | 1.89% | 2.96% | 4 | 21 | 34 |
| | Polymicrogyria | HP:0002126 | 2 | 1.38566e-6 | 9.24650e-3 | 4.62325e-3 | 3.0486 | 8.2005 | 25 | 1012 | 2.25% | 2.47% | 10 | 39 | 67 |
| | Uplifted earlobe | HP:0009909 | 3 | 9.47504e-6 | 6.32270e-2 | 2.10757e-2 | 3.4017 | 5.2914 | 18 | 653 | 1.62% | 2.76% | 2 | 3 | 4 |
| | Protuberant abdomen | HP:0001538 | 4 | 1.18974e-5 | 7.93916e-2 | 1.98479e-2 | 5.7667 | 1.7341 | 10 | 214 | 0.90% | 4.67% | 2 | 12 | 17 |
| | Pulmonic stenosis | HP:0001642 | 5 | 1.53953e-5 | 1.02733e-1 | 2.05465e-2 | 2.4379 | 11.8955 | 29 | 1468 | 2.61% | 1.98% | 11 | 38 | 55 |
| | Abnormality of the | HP:0002518 | 6 | 1.58370e-5 | 1.05681e-1 | 1.76134e-2 | 3.7701 | 3.9787 | 15 | 491 | 1.35% | 3.05% | 4 | 17 | 29 |

| | | | | | | | | | | | | | | | |
|---|---|---|---|---|---|---|---|---|---|---|---|---|---|---|---|
| | periventricular white matter | | | | | | | | | | | | | | |
| | Excessive salivation | HP:0003781 | 8 | 2.49276e-5 | 1.66342e-1 | 2.07927e-2 | 2.7704 | 7.9412 | 22 | 980 | 1.98% | 2.24% | 6 | 18 | 33 |
| | Abnormal pulmonary valve morphology | HP:0001641 | 9 | 4.27651e-5 | 2.85372e-1 | 3.17080e-2 | 2.2644 | 13.2488 | 30 | 1635 | 2.71% | 1.83% | 12 | 43 | 63 |
| | Severe muscular hypotonia | HP:0006829 | 10 | 4.43036e-5 | 2.95638e-1 | 2.95638e-2 | 3.8658 | 3.3628 | 13 | 415 | 1.17% | 3.13% | 5 | 27 | 39 |
| | Abnormality of the fingernails | HP:0001231 | 14 | 6.28269e-5 | 4.19244e-1 | 2.99460e-2 | 3.5187 | 3.9787 | 14 | 491 | 1.26% | 2.85% | 5 | 8 | 13 |
| | Drooling | HP:0002307 | 16 | 7.05757e-5 | 4.70951e-1 | 2.94345e-2 | 2.7272 | 7.3334 | 20 | 905 | 1.80% | 2.21% | 5 | 15 | 29 |
| | Cone-shaped epiphysis | HP:0010579 | 17 | 7.40366e-5 | 4.94046e-1 | 2.90615e-2 | 2.5710 | 8.5570 | 22 | 1056 | 1.98% | 2.08% | 6 | 21 | 29 |
| | Absent in utero rib ossification | HP:0006615 | 18 | 8.24000e-5 | 5.49855e-1 | 3.05475e-2 | 6.8560 | 1.0210 | 7 | 126 | 0.63% | 5.56% | 1 | 1 | 1 |
| | Absent in utero ossification of vertebral bodies | HP:0008435 | 18 | 8.24000e-5 | 5.49855e-1 | 3.05475e-2 | 6.8560 | 1.0210 | 7 | 126 | 0.63% | 5.56% | 1 | 1 | 1 |
| | Increased nuchal translucency | HP:0010880 | 18 | 8.24000e-5 | 5.49855e-1 | 3.05475e-2 | 6.8560 | 1.0210 | 7 | 126 | 0.63% | 5.56% | 1 | 1 | 2 |
| | Unossified sacrum | HP:0030290 | 18 | 8.24000e-5 | 5.49855e-1 | 3.05475e-2 | 6.8560 | 1.0210 | 7 | 126 | 0.63% | 5.56% | 1 | 1 | 1 |
| | Abnormal liver lobulation | HP:0100752 | 18 | 8.24000e-5 | 5.49855e-1 | 3.05475e-2 | 6.8560 | 1.0210 | 7 | 126 | 0.63% | 5.56% | 1 | 1 | 1 |
| | Broad forehead | HP:0000337 | 23 | 8.38995e-5 | 5.59861e-1 | 2.43418e-2 | 2.0148 | 17.8676 | 36 | 2205 | 3.25% | 1.63% | 8 | 35 | 49 |
| | Rod-cone dystrophy | HP:0000510 | 24 | 8.68398e-5 | 5.79482e-1 | 2.41451e-2 | 2.1432 | 14.4643 | 31 | 1785 | 2.80% | 1.74% | 12 | 79 | 127 |
| | Abnormal rib ossification | HP:0012306 | 26 | 1.05185e-4 | 7.01897e-1 | 2.69960e-2 | 6.5943 | 1.0615 | 7 | 131 | 0.63% | 5.34% | 1 | 2 | 5 |
| Mouse Phenotype Single KO | ostium primum atrial septal defect | MP:0010404 | 1 | 2.02072e-6 | 1.85037e-2 | 1.85037e-2 | 2.9837 | 8.3787 | 25 | 1034 | 2.25% | 2.42% | 7 | 16 | 17 |
| | abnormal Meibomian gland morphology | MP:0005252 | 2 | 4.47180e-6 | 4.09483e-2 | 2.04742e-2 | 4.1975 | 3.5735 | 15 | 441 | 1.35% | 3.40% | 3 | 12 | 16 |
| | abnormal skin sebaceous gland morphology | MP:0009535 | 3 | 5.54204e-6 | 5.07484e-2 | 1.69161e-2 | 4.1227 | 3.6383 | 15 | 449 | 1.35% | 3.34% | 3 | 13 | 18 |
| | ectopic ureter | MP:0011486 | 4 | 9.31434e-6 | 8.52914e-2 | 2.13228e-2 | 4.1935 | 3.3385 | 14 | 412 | 1.26% | 3.40% | 2 | 3 | 3 |
| | abnormal pericardial cavity morphology | MP:0012501 | 6 | 1.32280e-5 | 1.21129e-1 | 2.01881e-2 | 2.0617 | 20.3715 | 42 | 2514 | 3.79% | 1.67% | 22 | 91 | 118 |

| | | | | | | | | | | | | | | | |
|---|---|---|---|---|---|---|---|---|---|---|---|---|---|---|---|
| | pericardial edema | MP:0001787 | 7 | 1.45641e-5 | 1.33364e-1 | 1.90520e-2 | 2.2571 | 15.0639 | 34 | 1859 | 3.07% | 1.83% | 16 | 58 | 76 |
| | abnormal interatrial septum morphology | MP:0000282 | 8 | 1.61802e-5 | 1.48162e-1 | 1.85203e-2 | 2.0863 | 19.1722 | 40 | 2366 | 3.61% | 1.69% | 15 | 77 | 96 |
| | atrial septal defect | MP:0010403 | 10 | 1.83702e-5 | 1.68216e-1 | 1.68216e-2 | 2.1210 | 17.9162 | 38 | 2211 | 3.43% | 1.72% | 14 | 66 | 82 |
| | decreased cornea thickness | MP:0005543 | 11 | 3.43532e-5 | 3.14572e-1 | 2.85975e-2 | 3.7235 | 3.7599 | 14 | 464 | 1.26% | 3.02% | 4 | 15 | 17 |
| | abnormal prostate gland anterior lobe morphology | MP:0001163 | 13 | 4.43036e-5 | 4.05689e-1 | 3.12068e-2 | 3.8658 | 3.3628 | 13 | 415 | 1.17% | 3.13% | 3 | 13 | 14 |
| | abnormal brain ependyma motile cilium morphology | MP:0011056 | 14 | 5.23538e-5 | 4.79404e-1 | 3.42432e-2 | 5.4179 | 1.6612 | 9 | 205 | 0.81% | 4.39% | 4 | 10 | 16 |
| | abnormal ependyma motile cilium morphology | MP:0011059 | 14 | 5.23538e-5 | 4.79404e-1 | 3.42432e-2 | 5.4179 | 1.6612 | 9 | 205 | 0.81% | 4.39% | 4 | 10 | 18 |
| | absent Meibomian glands | MP:0006236 | 16 | 5.40483e-5 | 4.94921e-1 | 3.09325e-2 | 9.1413 | 0.6564 | 6 | 81 | 0.54% | 7.41% | 2 | 5 | 8 |
| | atrioventricular septal defect | MP:0010412 | 17 | 5.91320e-5 | 5.41472e-1 | 3.18513e-2 | 2.1898 | 14.1563 | 31 | 1747 | 2.80% | 1.77% | 9 | 55 | 75 |
| | abnormal head mesenchyme morphology | MP:0011260 | 18 | 6.18881e-5 | 5.66709e-1 | 3.14839e-2 | 3.1899 | 5.0159 | 16 | 619 | 1.44% | 2.58% | 5 | 22 | 28 |
| | absent subcutaneous adipose tissue | MP:0008843 | 19 | 9.01103e-5 | 8.25140e-1 | 4.34284e-2 | 3.8465 | 3.1197 | 12 | 385 | 1.08% | 3.12% | 3 | 15 | 19 |
| | abnormal kidney interstitium morphology | MP:0011425 | 20 | 9.72795e-5 | 8.90788e-1 | 4.45394e-2 | 2.3117 | 11.2473 | 26 | 1388 | 2.34% | 1.87% | 11 | 29 | 49 |
| | abnormal subcommissural organ morphology | MP:0009716 | 22 | 1.03657e-4 | 9.49191e-1 | 4.31450e-2 | 8.1368 | 0.7374 | 6 | 91 | 0.54% | 6.59% | 2 | 4 | 4 |
| | absent tracheal cartilage rings | MP:0004553 | 24 | 1.16811e-4 | 1.00000 | 4.45681e-2 | 3.3161 | 4.2218 | 14 | 521 | 1.26% | 2.69% | 2 | 3 | 3 |
| | abnormal atrioventricular septum morphology | MP:0010592 | 25 | 1.33537e-4 | 1.00000 | 4.89119e-2 | 2.0632 | 15.5096 | 32 | 1914 | 2.89% | 1.67% | 10 | 58 | 78 |
| Mouse Phenotype | small interparietal bone | MP:0004384 | 1 | 9.73658e-8 | 9.30914e-4 | 9.30914e-4 | 3.0878 | 9.7158 | 30 | 1199 | 2.71% | 2.50% | 10 | 21 | 24 |

| | | | | | | | | | | | | | | | |
|---|---|---|---|---|---|---|---|---|---|---|---|---|---|---|---|
| | complete atrioventricular septal defect | MP:0010413 | 2 | 3.34438e-7 | 3.19756e-3 | 1.59878e-3 | 3.6394 | 6.0450 | 22 | 746 | 1.98% | 2.95% | 4 | 20 | 30 |
| | abnormal vertebral body development | MP:0005227 | 3 | 4.56862e-7 | 4.36806e-3 | 1.45602e-3 | 4.4637 | 3.8085 | 17 | 470 | 1.53% | 3.62% | 3 | 10 | 12 |
| | pericardial edema | MP:0001787 | 4 | 2.09009e-6 | 1.99833e-2 | 4.99584e-3 | 2.0113 | 26.3517 | 53 | 3252 | 4.78% | 1.63% | 23 | 77 | 102 |
| | abnormal ureterovesical junction morphology | MP:0011488 | 6 | 3.07914e-5 | 2.94396e-1 | 4.90661e-2 | 3.2375 | 5.2509 | 17 | 648 | 1.53% | 2.62% | 3 | 5 | 6 |
| | abnormal intraocular pressure | MP:0005257 | 7 | 3.96052e-5 | 3.78665e-1 | 5.40950e-2 | 2.6160 | 8.7920 | 23 | 1085 | 2.07% | 2.12% | 9 | 23 | 26 |
| | increased hepatocyte apoptosis | MP:0003887 | 8 | 4.54044e-5 | 4.34111e-1 | 5.42639e-2 | 2.8175 | 7.0984 | 20 | 876 | 1.80% | 2.28% | 8 | 49 | 69 |
| | pancreas cysts | MP:0003336 | 9 | 4.55985e-5 | 4.35968e-1 | 4.84409e-2 | 3.2745 | 4.8862 | 16 | 603 | 1.44% | 2.65% | 5 | 13 | 21 |
| | absent Meibomian glands | MP:0006236 | 10 | 5.40483e-5 | 5.16756e-1 | 5.16756e-2 | 9.1413 | 0.6564 | 6 | 81 | 0.54% | 7.41% | 2 | 5 | 8 |
| | ostium primum atrial septal defect | MP:0010404 | 11 | 5.46298e-5 | 5.22316e-1 | 4.74832e-2 | 2.4466 | 10.2182 | 25 | 1261 | 2.25% | 1.98% | 7 | 22 | 23 |
| | fused kidneys | MP:0003605 | 12 | 5.73572e-5 | 5.48392e-1 | 4.56993e-2 | 4.0351 | 2.9739 | 12 | 367 | 1.08% | 3.27% | 4 | 6 | 6 |
| | common atrioventricular valve | MP:0010607 | 13 | 6.01714e-5 | 5.75299e-1 | 4.42538e-2 | 2.6800 | 7.8358 | 21 | 967 | 1.89% | 2.17% | 6 | 16 | 18 |
| | abnormal canal of Schlemm morphology | MP:0005204 | 14 | 6.11573e-5 | 5.84725e-1 | 4.17661e-2 | 2.8456 | 6.6771 | 19 | 824 | 1.71% | 2.31% | 6 | 13 | 14 |
| | macrophthalmia | MP:0001296 | 15 | 7.62318e-5 | 7.28852e-1 | 4.85902e-2 | 3.0056 | 5.6561 | 17 | 698 | 1.53% | 2.44% | 5 | 14 | 18 |
| | small zygomatic bone | MP:0004468 | 16 | 9.34700e-5 | 8.93667e-1 | 5.58542e-2 | 2.9548 | 5.7533 | 17 | 710 | 1.53% | 2.39% | 7 | 9 | 10 |
| | abnormal head mesenchyme morphology | MP:0011260 | 17 | 9.99293e-5 | 9.55424e-1 | 5.62014e-2 | 2.3533 | 10.6233 | 25 | 1311 | 2.25% | 1.91% | 11 | 37 | 45 |
| | abnormal subcommissural organ morphology | MP:0009716 | 18 | 1.03657e-4 | 9.91068e-1 | 5.50594e-2 | 8.1368 | 0.7374 | 6 | 91 | 0.54% | 6.59% | 2 | 4 | 4 |
| | exocrine pancreas atrophy | MP:0009164 | 19 | 1.14097e-4 | 1.00000 | 5.74149e-2 | 4.0401 | 2.7227 | 11 | 336 | 0.99% | 3.27% | 4 | 7 | 8 |
| | increased cardiomyocyte apoptosis | MP:0003222 | 21 | 1.35130e-4 | 1.00000 | 6.15228e-2 | 2.4637 | 8.9298 | 22 | 1102 | 1.98% | 2.00% | 9 | 52 | 60 |
| | small vertebral body | MP:0004670 | 22 | 1.40711e-4 | 1.00000 | 6.11516e-2 | 2.8543 | 5.9559 | 17 | 735 | 1.53% | 2.31% | 3 | 14 | 17 |

| | |
|---|---|
| The test set contains 1,109 (1%) of all 136,859 regions. | The test set picked 1,072 genes, the background set picked 11,393 genes. |
| *Ensembl Genes* has 18,777 terms covering 18,777 (100%) of all 18,777 genes. | 18,777 ontology terms were tested (100%) using an annotation count range of [1, 1000]. |
| *GO Biological Process* has 13,159 terms covering 16,804 (89%) of all 18,777 genes. | 13,159 ontology terms were tested (100%) using an annotation count range of [1, 1000]. |
| *GO Cellular Component* has 1,728 terms covering 17,911 (95%) of all 18,777 genes. | 1,728 ontology terms were tested (100%) using an annotation count range of [1, 1000]. |
| *GO Molecular Function* has 4,222 terms covering 16,729 (89%) of all 18,777 genes. | 4,222 ontology terms were tested (100%) using an annotation count range of [1, 1000]. |
| *Human Phenotype* has 6,673 terms covering 3,390 (18%) of all 18,777 genes. | 6,673 ontology terms were tested (100%) using an annotation count range of [1, 1000]. |
| *Mouse Phenotype Single KO* has 9,157 terms covering 9,525 (51%) of all 18,777 genes. | 9,157 ontology terms were tested (100%) using an annotation count range of [1, 1000]. |
| *Mouse Phenotype* has 9,561 terms covering 9,709 (52%) of all 18,777 genes. | 9,561 ontology terms were tested (100%) using an annotation count range of [1, 1000]. |

| |
|---|
| GREAT version 4.0.4 |
| Species assembly: hg38 |
| Association rule: Basal+extension: 5000 bp upstream, 1000 bp downstream, 1000000 bp max extension, curated regulatory domains included |
